# TreeCluster: clustering biological sequences using phylogenetic trees

**DOI:** 10.1101/591388

**Authors:** Metin Balaban, Niema Moshiri, Uyen Mai, Siavash Mirarab

## Abstract

Clustering homologous sequences based on their similarity is a problem that appears in many bioinformatics applications. The fact that sequences cluster is ultimately the result of their phylogenetic relationships. Despite this observation and the natural ways in which a tree can define clusters, most applications of sequence clustering do not use a phylogenetic tree and instead operate on pairwise sequence distances. Due to advances in large-scale phylogenetic inference, we argue that tree-based clustering is under-utilized. We define a family of optimization problems that, given a (not necessarily ultrametric) tree, return the minimum number of clusters such that all clusters adhere to constraints on their heterogeneity. We study three specific constraints that limit the diameter of each cluster, the sum of its branch lengths, or chains of pairwise distances. These three versions of the problem can be solved in time that increases linearly with the size of the tree, a fact that has been known by computer scientists for two of these three criteria for decades. We implement these algorithms in a tool called TreeCluster, which we test on three applications: OTU picking for microbiome data, HIV transmission clustering, and divide-and-conquer multiple sequence alignment. We show that, by using tree-based distances, TreeCluster generates more internally consistent clusters than alternatives and improves the effectiveness of downstream applications. TreeCluster is available at https://github.com/niemasd/TreeCluster.

## Introduction

Homologous molecular sequences across different species or even within the same genome can show remarkable similarity due to their shared evolutionary history. These similarities have motivated many applications to first group the elements of a diverse set of sequences into *clusters* of set of sequences with high similarity for use in subsequent steps. The precise meaning of clusters depends on the application. For example, when analyzing 16S microbiome data, the standard pipeline is to use Operational Taxonomic Units (OTUs), which are essentially clusters of closely related sequences that do not diverge more than a certain threshold [1–3]. Another example is HIV transmission inference, a field in which a dominant approach is to cluster HIV sequences from different individuals based on their similarity (again using a threshold) and to use these clusters as proxies to clusters of disease transmission [4, 5].

Shared evolutionary histories, the origin of similarity among homologous sequences, can be captured by a phylogenetic tree. The true phylogeny is never known, but it can be inferred from sequence data, [6, 7] and recent advances have led to methods that can infer approximate maximum-likelihood (ML) phylogenetic trees in sub-quadratic time, which can easily scale to datasets of even millions of sequences [8]. Moreover, accurate alignment of datasets with hundreds of thousands of species (a prerequisite to most phylogenetic reconstruction methods) is now possible using divide-and-conquer methods [9, 10].

Most existing sequence clustering methods use the pairwise distances among sequences as input but do not take advantage of phylogenetic trees. For example, the widely-used UCLUST [2] searches for a clustering that minimizes the Hamming distance of sequences to the cluster centroid while maximizes the Hamming distance between centroids. Several other clustering methods have been developed for various contexts, such as gene family circumscription [11, 12] and large protein sequence databases [13].

Using phylogenies for clustering has two potential advantages. *i*) Since phylogenies explicitly seek to infer the evolutionary history, phylogeny-based clustering has the potential to not only reflect evolutionary distances (i.e., branch lengths) but also relationships (i.e., the tree topology). Recall also that branch lengths in a phylogeny are model-based “corrections” of sequence distances in a statistically-rigorous way [7], and therefore, may better reflect divergence between organisms. *ii*) When inferred using subquadratic algorithms, the tree can eliminate the need to compute all pairwise distances, which can improve speed and scalability. Moreover, a phylogeny often has to be inferred for purposes other than clustering and thus typically is readily available. However, despite these potentials, to our knowledge, no systematic method for phylogeny-guided clustering exists. Built for analyzing HIV transmissions, ClusterPicker [14] clusters sequences based on their distances while using the phylogenetic tree as a constraint; however, it still uses sequence (not tree) distances and scales cubically with respect to number of sequences in the worst case.

Given a rooted phylogenetic tree, if the tree is ultrametric (that is, distances of all the leaves to the root are identical), clustering sequences based on the tree can proceed in an obvious fashion: the tree can be cut at some distance from the root, thereby partitioning the tree into clusters (Fig. 1a). This approach extends in natural ways to unrooted ultrametric trees by first rooting the tree at the unique midpoint [15] and proceeding as before. However, inferred phylogenetic trees are rarely ultrametric. Different organisms can evolve with different rates of evolution, and even when the rates are identical (leading to an ultrametric true tree), there is no guarantee that the inferred trees will be ultrametric. Given a non-ultrametric (and perhaps unrooted) tree, the best way to cluster sequences is not obvious (Fig. 1b).

**Fig 1.**
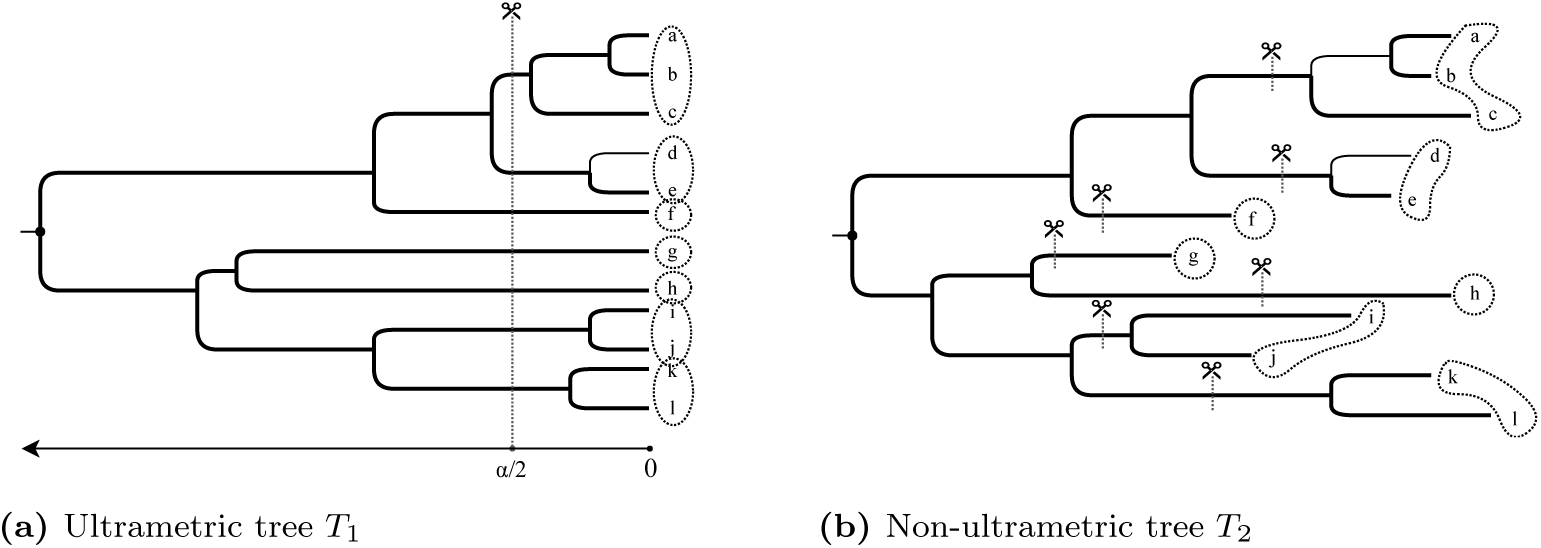
When the phylogenetic tree is ultrametric, clustering is trivial: for a threshold *α*, cut the tree at 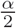 height (a) When the tree is not ultrametric, it is not obvious how to cluster leaves (b). In either cases, a set of cut edges defines a clustering.

One way to approach tree-based clustering is to treat it as an optimization problem. We can define problems of the following form: “find the minimum number of clusters such that some criteria constrain each cluster.” Interestingly, at least two forms of such optimization problems have been addressed as early as the 1970s by the theoretical computer science community, in the context of proving more challenging theorems. *Tree partitioning* problems were defined to partition any arbitrary tree into minimum number of subtrees such that the maximum path length between any nodes [16] or sum of all node weights [17] in each subtree are constrained by a given threshold. Both problems can be solved exactly using straightforward linear-time algorithms; these algorithms, however, to our knowledge, are mostly ignored by bioinformaticians.^1^

Here, we argue that the tree-based clustering approach should be revitalized in the field of bioinformatics. In this paper, we introduce a family of tree partitioning problems and describe linear-time solutions for three instances of the problem (two of which correspond to the aforementioned max and sum problems with known algorithms). More importantly, we show that these tree-based clustering algorithms can result in improved downstream biological analyses in three different contexts: defining microbial OTUs, HIV transmission clustering, and divide-and-conquer multiple sequence alignment.

## Materials and methods

### TreeCluster

#### Problem definition

Let *T* = (*V, E*) be an unrooted binary tree represented by an undirected acyclic graph with vertices *V* either degree one or three, weighted edges *E*, and leafset ℒ. We denote the path length between leaves *u* and *v* on *T* with *d*_*T*_ (*u, v*) or simply *d*(*u, v*) when clear by context. The weight of an edge (*u, v*) (i.e., its branch length) is denoted by *w*(*u, v*). A natural way to define a clustering of the leaves in ℒ is to associate a clustering to a cut set *C* ⊆*E* on the edges of *T*. We define a partition {*L*_1_, *L*_2_, … *L*_*N*_} of ℒ to be an *admissible* clustering if it can be obtained by removing some edge set *C* from *E* and assigning leaves of each of the resulting connected components to a set *L*_*i*_ (note: *N* ≤ |*C*| + 1).

For a given tree *T*, let *f*_*T*_: 2^ℒ^ → ℝ be a function that maps a subset of the leafset ℒ to a real number. The purpose of *f*_*T*_ (.) is to characterize the diversity of elements at the leaves within each cluster, and it is often defined as a function of the edge weights in the cluster. For example, it can be the diameter of a subset: *f*_*T*_ (*L*) = max_*u,v*∈*L*_ *d*_*T*_ (*u, v*). We define a family of problems that seek to minimize the number of clusters while each cluster has to adhere to constraints defined using *f*_*T*_ (.). More formally:

#### Definition 1

(Min-cut partitioning problem family). *Given a tree T with leafset* ℒ *and a real number α, find an admissible partition* {*L*_1_ … *L*_*N*_*} of* ℒ *that satisfies* ∀*i, f*_*T*_ (*L*_*i*_)≤*α and has the minimum possible cardinality (N) among all such clusterings*.

A natural way to limit the diversity within a cluster is to constrain all pairwise distances among members of the cluster to be less than a given threshold:

#### Definition 2

(Max-diameter min-cut partitioning problem). *The Min-cut partitioning problem (Definition 1) is called Max-diameter min-cut partitioning problem when* 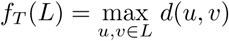.

One potential disadvantage of max diameter min-cut partitioning is its susceptibility to outliers: the largest distance within a cluster may not be always an accurate representation of the degree of diversity in the cluster. A natural choice that may confine the effect of outliers is the following:

#### Definition 3

(Sum-length min-cut partitioning problem). *The Min-cut partitioning problem is called Sum-length min-cut partitioning problem when* 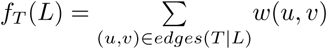 *where T |L is the tree T restricted to a subset of leaves L*.

We also study a fourth problem, which we will motivate later:

#### Definition 4

(Single-linkage min-cut partitioning problem). *The Min-cut partitioning problem is called Single-linkage min-cut partitioning problem when* 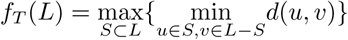.

Next, we will show linear-time algorithms for the Max-diameter, Sum-length, and Single-linkage min-cut partitioning problems. Note that all of these algorithms use variations of the same greedy algorithm. Two of these greedy algorithms (max and sum) are already described in the theoretical computer science literature. Nevertheless, we reiterate the solutions using consistent terminology and provide alternative proofs of their correctness.

### Linear-time solution for Max-diameter min-cut partitioning

A linear-time solution for the Max-diameter min-cut partitioning problem was first published by Parley *et al*. [16]. We present Algorithm 1, which is similar to the algorithm by Parley *et al*. (adding branch lengths), and we give an alternative proof.

The algorithm operates on *T*′, which is an arbitrary rooting of *T* at node *r*. We denote the subtree rooted at an internal node *u* as *U*. Let the two children of *u* be called *u*_*l*_ and *u*_*r*_, and let the tree rooted by them be *U*_*l*_ and *U*_*r*_. We use *w*_*l*_ and *w*_*r*_ to denote *w*(*u, u*_*l*_) and *w*(*u, u*_*r*_), respectively. We define *B*(*u*) to be the length of the path from *u* to the farthest *connected* leaf in *U* under the optimal clustering.

The algorithm uses a bottom-up traversal of the tree. When we arrive at node *u*, one or more new paths form between the two trees *U*_*r*_ and *U*_*l*_. Among those paths, the longest one has the length *B*(*u*_*l*_) + *w*_*l*_ + *B*(*u*_*r*_) + *w*_*r*_. If this value exceeds the threshold, we break either (*u, u*_*r*_) or (*u, u*_*l*_), depending on which minimizes *B*(*u*). Note that the algorithm always cuts at most one child edge of every node, and thus, *B*(*u*) is always well-defined.

We now show the algorithm correct. Let *A*(*u*) be the minimum number of clusters under *U*, each with a diameter less than *α*; i.e., *A*(*r*) is the objective function.

#### Theorem 1

*Algorithm 1 computes a clustering with minimum A*(*r*) *for rooted tree T*′. *In addition, among all possible such clusterings, the algorithm picks the solution with minimum B*(*r*).

*Proof*. The proof uses induction. The base case for the induction is the simple rooted tree with root *u* and two leaves *u*_*l*_ and *u*_*r*_. If *w*_*l*_ + *w*_*r*_ *> α* the algorithm cuts the longer branch whereas if *w*_*l*_ + *w*_*r*_ ≤ *α* no branch is cut. In both cases, the theorem holds.

Th inductive hypothesis is that for a node *u*, the algorithm has computed *A*(*u*_*l*_), *A*(*u*_*r*_), *B*(*u*_*l*_), and *B*(*u*_*r*_) optimally. We need to prove that a solution other than the one computed by our algorithm *i*) cannot have a lower number of clusters, call it *A*′(*u*), and *ii*) when *A*′(*u*) = *A*(*u*), cannot have a lower distance to the farthest connected leaf, call it *B*′(*u*).

#### Algorithm 1: Linear solution for Max diameter min-cut partitioning

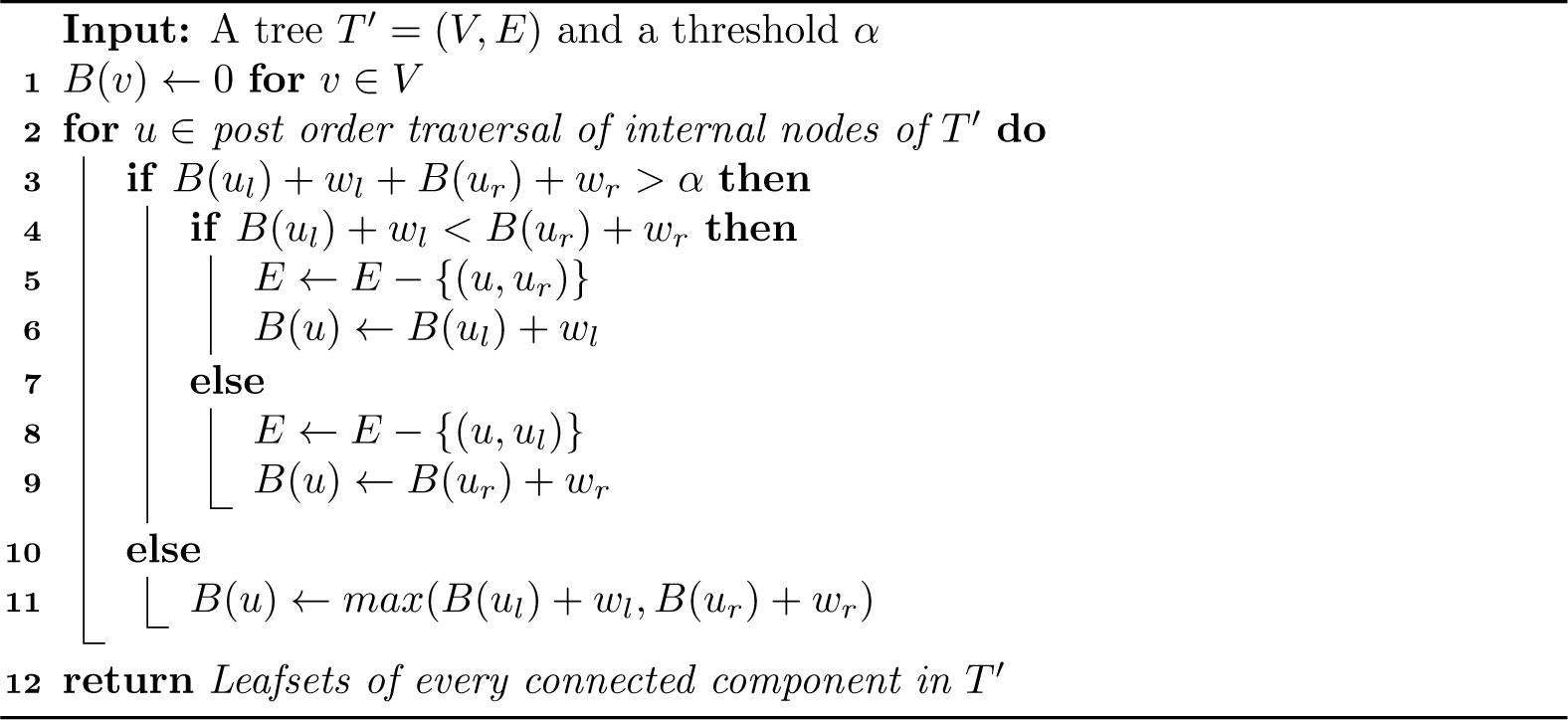

When *B*(*u*_*l*_) + *w*_*l*_ + *B*(*u*_*r*_) + *w*_*r*_≤*α*, we have *A*(*u*) = *A*(*u*_*l*_) + *A*(*u*_*r*_) −1, which is the minimum possible by inductive hypothesis and the fact that the number of clusters cannot go down by more than one on node *u*. Also, *B*(*u*) is optimal by construction.

When *B*(*u*_*l*_) + *w*_*l*_ + *B*(*u*_*r*_) + *w*_*r*_ *> α*, without loss of generality, assume that *B*(*u*_*l*_) + *w*_*l*_ ≥*B*(*u*_*r*_) + *w*_*r*_ and thus, the algorithm cuts the (*u, u*_*l*_) branch, getting *A*(*u*) = *A*(*u*_*l*_) + *A*(*u*_*r*_) and *B*(*u*) = *B*(*u*_*r*_) + *w*_*r*_. Note that *A*′(*u*) *< A*(*u*) is only possible if *A*′(*u*_*l*_) = *A*(*u*_*l*_) and *A*′(*u*_*r*_) = *A*(*u*_*r*_) and we do not cut any branch at *u* in the alternative clustering. However, this scenario is *not* possible because

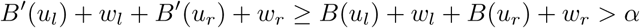

where the first inequality follows from the inductive hypothesis and the final inequality shows that we will have to cut a branch in any alternative setting. Finally, we need to show that an alternative solution with *A*′(*u*) = *A*(*u*) but *B*′(*u*) *< B*(*u*) is not possible. The inequality requires that either *B*′(*u*_*l*_) *< B*(*u*_*l*_) or *B*′(*u*_*r*_) *< B*(*u*_*r*_). First, consider the *B*′(*u*_*l*_) *< B*(*u*_*l*_) case, which is possible only if *A*′(*u*_*l*_) = *A*(*u*_*l*_) + 1. Note that *A*′(*u*) = *A*(*u*) requires *A*′(*u*_*r*_) = *A*(*u*_*r*_) (and thus *B*′(*u*_*r*_) = *B*(*u*_*r*_)) and that *B*′(*u*_*l*_) + *w*_*l*_ + *B*(*u*_*r*_) + *w*_*r*_ *< α*, which is possible. Under this condition, we find:

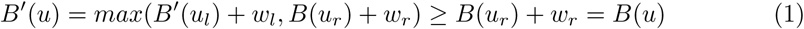

If instead *B*′(*u*_*r*_) *< B*(*u*_*r*_), similar conditions can be written, resulting in

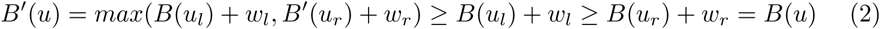

Thus, *A*(*u*) and *B*(*u*) are optimal when *B*(*u*_*l*_) + *w*_*l*_ + *B*(*u*_*r*_) + *w*_*r*_ *> α*.

#### Corollary 1

*Let C*′ *be the cut set obtained by running Alg. 1 on any arbitrary rooting T*′ *of unrooted tree T. C*′ *optimally solves the Max-diameter min-cut partitioning problem*.

*Proof*. Let *r*_*r*_ and *r*_*l*_ denote the right and the left child of the root of *T*′. Every edge in *T* can be mapped to *T*′ except the edge (*r*_*right*_, *r*_*left*_), from which we define a mapping to (*r, r*_*right*_) (w.l.o.g). Using this mapping, the optimal clustering (i.e. optimal cut-set) on *T* can be translated to an alternative max diameter min-cut partitioning on *T*′. However, by Theorem 1, *A*(*r*) is optimal and cannot be improved by any alternative partitioning. Since any admissible clustering on *T*′ is also admissible on *T*, Alg. 1 minimizes *N*.

### Linear solution for the Sum-length min-cut partitioning problem

A linear algorithm which partitions trees into the fewest clusters having total node weights lower than or equal to *α* has been previously published by Kundu *et al*. [17]. In order to solve Sum-length min-cut partitioning problem, we require an altered version of the original algorithm that works on edge (instead of node) weights and focuses on binary trees. Algorithm 1 with two simple modifications solves Sum-length min-cut partitioning problem optimally (see Alg. 3 in Appendix B). The first modification is that here we define the auxiliary variable *B*(*u*) denoting sum of weights of all descendent edges connected to *u* at the stage it is processed by the algorithm. Secondly, in the bottom-up traversal of internal nodes of *T*′, for node *u*, w.l.o.g, let *B*(*u*_*l*_) + *w*_*l*_ *> B*(*u*_*r*_) + *w*_*r*_. If the sum of branch lengths in the combined subtree exceed *α*, we break the edge (*u, u*_*l*_). Unlike Algorithm 1, where *B*(*u*_*l*_) + *w*_*l*_ + *B*(*u*_*r*_) + *w*_*r*_ ≤ *α*, here, *B*(*u*) is set to *B*(*u*_*l*_) + *w*_*l*_ + *B*(*u*_*r*_) + *w*_*r*_. The proof for the correctness of the algorithm is analogous to that of Alg. 1 and is given in Appendix B.

### Single linkage min-cut partitioning

We now address the Single-linkage problem (Definition 4), which can be considered a relaxation of the max diameter min-cut partitioning. To motivate this problem, first consider the following definition.

#### Definition 5

(Single-linkage clustering). *We call a partition of* ℒ *to be a Single-linkage clustering when for every a, b* ∈ ℒ, *a and b are in the same cluster if and only if there exist a chain* ℋ = *c*_0_, *c*_1_ …, *c*_*m*_, *c*_*m*+1_, *where a* = *c*_0_ *and b* = *c*_*m*+1_, *and for every* 0 ≤ *i* ≤ *m, we have d*(*c*_*i*_, *c*_*i*+1_) ≤ *α*.

Thus, every pair of nodes are put in the same cluster if (but not only if) their distance is below the threshold (the rest follows from transitivity). The next result (proved in Appendix B.) motivates the choice of *f*_*T*_ (.) in Definition 4.

#### Proposition 1

*The optimal solution to the Single-linkage min-cut partitioning problem (Definition 4) is identical to the Single-linkage clustering of Definition 5*.

We now present Algorithm 2, a linear-time solution to the Single-linkage min-cut partitioning problem. The algorithm first computes the distances to the closest node on left, right, and outside each node *u* in a post-order followed by a pre-order traversal. Then, on a post-order traversal, it cuts each child edge iff the minimum distance of leaves under it to leaves under its sibling *and* to any leaf outside the node both exceed the threshold.

#### Theorem 2

*A min-cut partitioning computed by Algorithm 2 optimally solves Single-linkage min-cut partitioning problem (Definition 5)*.

*Proof*. Let *a* ↭ *b* be the path between leaves *a* and *b* on *T*. Fixing *a* and *b*, for each node *j*, we use the term *support of j* denoted by *s*(*j*) to refer to the unique node on all the three paths *a* ↭ *b, a* ↭ *j*, and *b* ↭ *j*. We refer to a group of leaves that share a mutual support with respect to *a* and *b* as a *bubble* (e.g., triangles in Fig. 2). Among all bubbles branching out of *a* ↭ *b*, let the one with the closest support to *a* be *A*′. We name the leaf closest to *a* on *A*′ as *a*′ (Fig. 2).

**Fig 2.**
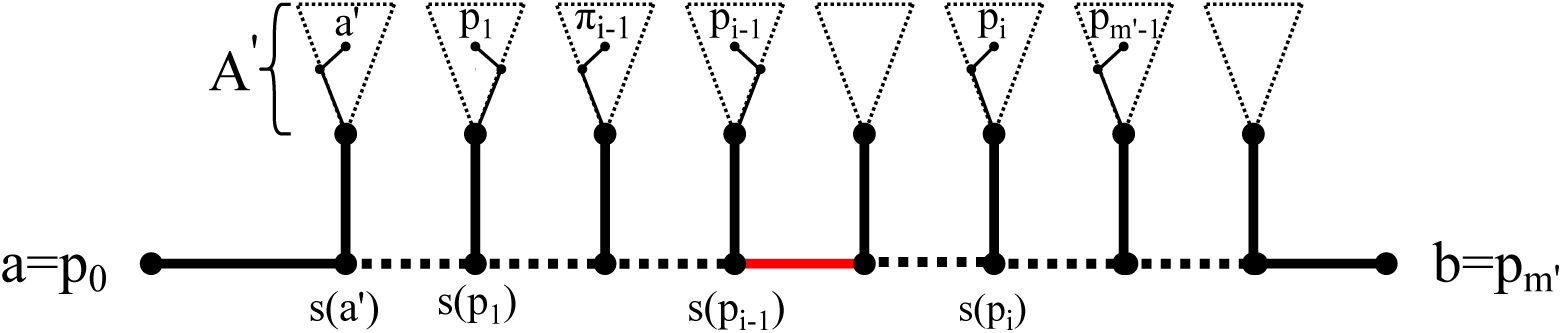
A sketch showing the setup for constructing the chain ℋ.

#### Algorithm 2: Single-Linkage Single-linkage min-cut partitioning

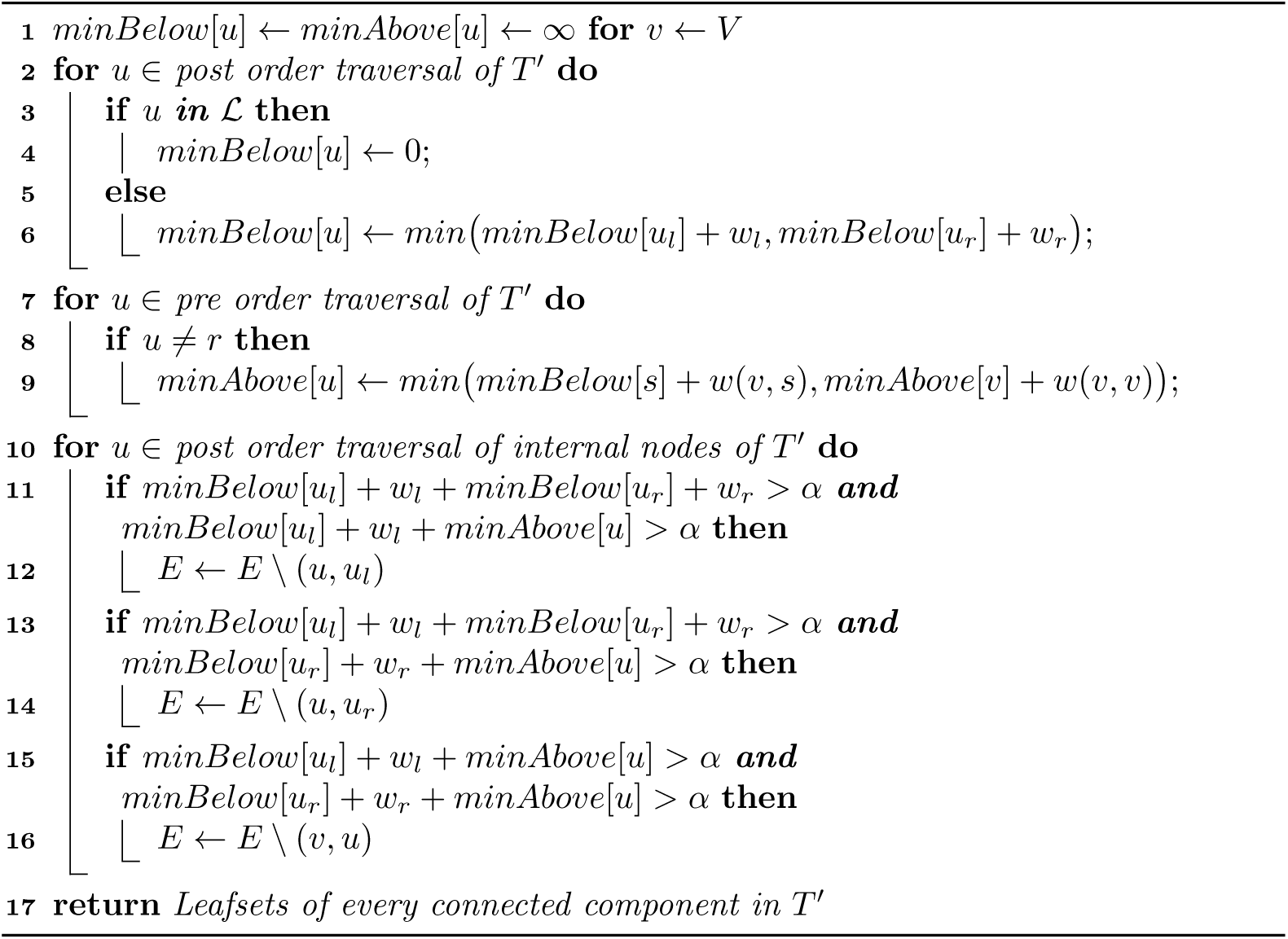

- If *d*(*a, b*) ≤ *α* holds, the algorithm will never cut any edge on *a* ↭*b* due to the following observation. For every internal node *u* on *a* ↭*b*, let *v* and *w* be the adjacent nodes on *a* ↭*u* and *u* ↭*b* respectively. Also let *p*_*a*_ be the closest leaf to *u* whose support *s*(*p*_*a*_) is on *a* ↭*u*, and *p*_*b*_ to be the closest leaf to *u* whose support *s*(*p*_*b*_) is on *u* ↭*b*. *d*(*p*_*a*_, *u*) + *d*(*u, p*_*b*_)≤ *d*(*a, u*) + *d*(*u, b*) ≤ *α* holds, therefore regardless of the rooting, (*v, u*) and (*u, w*) is never cut by Alg. 2.
- if a chain ℋ exists, due to the previous observation, there is no cuts on *c*_*i*_ ↭ *c*_*i*+1_ for every 0≤ *i* ≤ *m*. Consequently, *a* and *b* are connected through a path and hence are in the same cluster.
- Assume Alg. 2 places *a* and *b* on the same cluster, i.e. does not cut any edge on *a*↭ *b*. We present a procedure to generate a chain ℋ described in Definition 1. We define *p*_0_ = *a* and *p*_*m*_*I* = *b*. For 1 ≤ *i* ≤ *m*′, we let *p*_*i*_ be the closest leaf to *p*_*i-*1_

whose support *s*(*p*_*i*_) is on *p*_*i-*1_ ↭ *b* and *s*(*p*_*i*_) ≠*s*(*p*_*i-*1_) (i.e., *p*_*i*_ is in the bubble to the right of the bubble of *p*_*i-*1_). Conversely, for 1 ≤ *i* ≤ *m*′, we let *π*_*i*_ denote the closest leaf to *p*_*i*_ whose support is on *a* ↭ *s*(*p*_*i-*1_) (i.e., is in a bubble to the left of *p*_*i*_); note *π*_1_ = *a*. Every *π*_*i*_ ∈ {*p*_0_ … *p*_*i-*1_} due to following observation: if not, *s*(*π*_*i*_) has to be on *s*(*p*_*j-*1_) ↭ *s*(*p*_*j*_) for some *j*; but, we would have *d*(*p*_*j-*1_, *π*_*i*_) ≤ *d*(*p*_*j-*1_, *p*_*j*_), which contradicts the definition of *p*_*i*_. The fact that Alg. 2 retains (*a, s*(*a*′)) indicates that *min*(*d*(*a, a*′), *d*(*a, p*_1_)) = *d*(*a, p*_1_) ≤ *α*; therefore, we add *a* → *p*_1_ to an auxiliary graph ℋ ′. Now, consider Alg. 2 when it processes the node *s*(*p*_*i-*1_) for 1 *< i*. The fact that the first edge on path *s*(*p*_*i-*1_) ↭ *s*(*p*_*i*_) (shown in red color in Fig. 2) is not cut indicate that either *d*(*π*_*i-*1_, *p*_*i*_) ≤ *α* or *d*(*p*_*i-*1_, *p*_*i*_) ≤ *α*. Depending on which is true, we add a link from *π*_*i-*1_ → *p*_*i*_ or *p*_*i-*1_ → *p*_*i*_ to ℋ ′. We repeat this process for all *i* until we reach *i* = *m*′, where we add an edge to *p*_*m*_*′* = *b*. Noting that *π*_*i*_ ∈ {*p*_0_ … *p*_*i-*1_}, the ℋ′ graph becomes a directed tree, rooted at *a* with a directed path to the leaf *b*. This directed path constitute the valid chain ℋ.

### Clade constraint for rooted trees

So far, we have focused on unrooted trees. This choice is partially driven by the fact that phylogenetic reconstruction tools predominantly use time reversible models of sequence evolution (e.g., GTR [18]), and therefore output an unrooted tree. Nevertheless, researches have developed various methods for rooting trees [19, 20] including scalable algorithms [15]. When a rooted tree is available, each “monophyletic clade”, i.e. a group of entities that includes all descendants of their common ancestor, are biologically meaningful units. Thus, we may want to constrain each cluster to be a clade. Such “clade” constraints, in fact, make clustering easier; our algorithms can be easily altered to ascertain that each cluster is also a clade. Specifically on Algorithm 1, when we have *B*(*u*_*l*_) + *w*_*l*_ + *B*(*u*_*r*_) + *w*_*r*_ *> α*, we simply need to cut both (*u, u*_*l*_) and (*u, u*_*r*_) (instead of cutting only the longer one). This small modification allows Max-Diameter, Sum-length, and Single-linkage min-cut partitioning problems to be solved in linear time while imposing the clade constraint.

### Three Applications of TreeCluster

While sequence clustering has many applications, in this paper, we highlight three specific areas as prominent examples.

#### Application 1: OTU clustering

##### Biological Problem

For microbiome analyses using 16S sequences generated from whole communities, the standard pipeline uses operational taxonomic units (OTUs). Sequences with similarity at or above a certain threshold (say 97%) are grouped into OTUs, which are the most fine-grained level at which organisms are distinguished. All sequences assigned to the same OTU are treated as one organism in downstream analyses, such as taxonomic profiling, taxonomic identification, sample differentiation, or machine learning. The use of a similarity threshold instead of a biological concept of species is to avoid the notoriously difficult problem of defining species for microbial organisms. In addition, the use clusters of similar sequences as OTUs can provide a level of robustness with respect to sequencing errors.

Most applications of OTUs are closed-reference: a reference database of known organisms is selected and OTUs are defined for reference sequences using methods such as UCLUST [2] and Dotur [3]. These methods cluster sequences based on a chosen threshold of similarity, often picking a centroid sequence to represent an OTU. Reads from a 16S sample are then compared to the OTUs, and the closest OTU is found for each read (judging distance by sequence similarity). Once all reads are processed for all samples, an OTU table can be built where the rows are samples, the columns are OTUs, and each cell gives the frequency of an OTU in a sample. This table is then used in downstream analyses. Several large reference databases exist for these OTU-based analyses [21–23]. One of these databases, popularized through pipelines such as Qiita [24], is Greengenes [23].

Regardless of the downstream application of an OTU table, one would prefer the OTUs to be maximally coherent (i.e., internally consistent) so that they represent organisms as faithfully as possible. We will focus our experiments on the closed-reference OTU picking methods and the Greengenes as the reference library. However, note that open-reference OTU picking and sub-operational-taxonomic-unit (sOTU) methods [25–27] also exist and involve a similar need for sequence clustering.

##### Existing methods

Several hierarchical clustering tools have been proposed for OTU clustering [3, 28]. Non-hierarchical clustering methods [2, 29] however gained popularity due to being computationally less demanding compared to hierarchical methods. Two prominent methods are UCLUST [2] and CD-HIT [29], which share the same algorithmic strategy: for a given threshold *α*, UCLUST determines a set of representative sequences dynamically by assigning query sequences into representative sequences (centroids) such that, ideally, distance between each query and its assigned centroid is less than *α* while distances between centroids is more than *α*. UCLUST is a heuristic algorithm, and the processing order of the queries may affect the outcome clustering. CD-HIT is different from UCLUST mostly in its strategy for computing distances.

##### Formulation as min-cut partitioning

We define OTUs by solving the Min-diameter, Sum-Length, or Single-linkage min-cut partitioning problems using a chosen threshold *α* and an inferred ML phylogeny. Each cluster in the resulting partition is designated as an OTU.

##### Experiments

We evaluate the quality of tree-based OTU clustering by comparing it to UCLUST as used by Greengenes [23]. We run TreeCluster on the phylogenetic tree of 203,452 sequences in Greengenes v13.5 database in three modes: max, sum, and Single-linkage. We use the following 20 thresholds: [0.005, 0.05] with a step size of 0.005 and (0.05, 0.15] with a step size of 0.01. For Single-linkage, we only go up to 0.1 because above this threshold, the number of clusters becomes much smaller than other methods.

From the same Greengenes database, we extract OTU clusters for all available sequence identity thresholds up to 0.15 (i.e., 0.03, 0.06, 0.09, 0.12, and 0.15). We measure the quality of a clustering {*L*_1_, …, *L*_*N*_*}*by its weighted average of average pairwise distance per cluster (which we call *cluster diversity* for shorthand), given by the following formula:

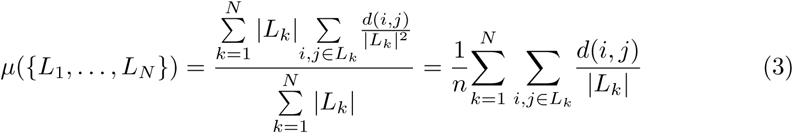

where *n* denotes the number of sequences clustered. We compute distance *d*(*i, j*) between two elements using two methods: *tree distance*, which is the path lengths on the inferred phylogenetic tree, or the sequence-based hamming distance. Hamming distances are computed pairwise from the multiple sequence alignment of all 203,452 sequences in Greengenes database and ignore any site that includes a gap in the pairwise alignment. Clearly, cluster diversity alone is not sufficient to judge results (singletons have zero diversity). Instead, we compare methods at the same level of clustering with respect to their diversity. Thus, as we change the threshold *α*, we compare methods for choices of the threshold where they result in (roughly) equal numbers of clusters. Given the same number of clusters, a method with lower cluster diversity is considered preferable.

#### Application 2: HIV transmission cluster analyses

##### Biological Problem

HIV sequences evolve fast and therefore their phylogenetic relationships have a trace of the transmission chains from one person to another [30]. The ability to perform phylogenetic analyses of HIV sequences is critical for epidemiologists who design and evaluate HIV control strategies [31–35]. The results of these analyses can provide information about the genetic linkage [36] and transmission histories [37], as well as mixing across subpopulations [38]. A recent advancement in computational molecular epidemiology is the use of transmission clustering to predict at-risk individuals and epidemic growth: infer transmission clusters from pairwise sequence distances, monitor the growth of clusters over time, and prioritize clusters with the highest growth rates [39]. In this monitoring framework, two natural questions come about: What is the optimal way to infer transmission clusters from molecular data, and how can transmission cluster inference be performed more efficiently?

##### Existing methods

Two existing tools perform such clustering. Cluster Picker [4] is given a distance threshold, a phylogenetic tree, and sequences. It clusters individuals such that each cluster defines the leaves of a clade in the tree, the maximum pairwise sequence-based distance in each cluster is below the threshold, and the number of clusters is minimized. HIV-TRACE is a tool that, given a distance threshold and sequences, clusters individuals such that, for each pair of individuals *u* and *v*, if the Tamura-Nei 93 (TN93) distance [40] between *u* and *v* is below the threshold, *u* and *v* are placed in the same cluster [5]. Both methods scale worse than linearly with the number of sequences (quadratically and cubically, respectively, for HIV-Trace and Cluster Picker), and for large datasets, they can take hours, or even days, to run (however, HIV-Trace does enjoy trivial parallelism).

##### Formulation as min-cut partitioning

Transmission clustering is similar to our problem formulation in that it involves cutting edges such that the resulting clusters (as defined by the leafsets resulting from the cuts) must adhere to certain constraints. Both Cluster Picker and HIV-TRACE utilize pairwise distances computed from sequences, but when reformulated to utilize tree-based distances from an inferred phylogeny, Cluster Picker becomes analogous to our Max-diameter min-cut partitioning (with an added constraint that clusters must define clades in the phylogeny), and HIV-TRACE analogous to the Single-linkage min-cut partitioning.

##### Experiments

To evaluate the effectiveness of HIV transmission clustering, we first simulate HIV epidemic data using FAVITES [41]. For the simulation parameters, we use the parameters described in Moshiri *et. al*. [41] to model the San Diego HIV epidemic between 2005 and 2014. However, we deviate from the original parameter set in one key way: originally, all HIV patients were sequenced at the end time of the epidemic, yielding an ultrametric tree in the unit of time, but to better capture reality, we instead sequence each patient the first time they receive Antiretroviral Treatment (ART). In our simulations, we vary two parameters: the expected time to begin ART as well as the expected degree of the social contact network, which underlies the transmission network.

Higher ART rates and lower degrees both result in a slower epidemic and change patterns of phylogenetic branch length [41]. The complete FAVITES parameter set can be found in the supplementary materials. We infer phylogenies from simulated sequences under the GTR+Γ model using FastTree-II [8], and we use MinVar algorithm to root the trees using FastRoot [42].

We use HIV-TRACE [5] as well as multiple clustering modes of TreeCluster to infer transmission clusters. We were unable to use Cluster Picker [4] due to its excessive running time. For HIV-TRACE, we use a clustering threshold of 1.5% as suggested by its authors [39]. Because HIV-TRACE estimates pairwise sequences distances under the TN93 model, [40] which tend to be underestimates of phylogenetic distance estimated under the GTR model, we use a clustering threshold of 3% for Single-Linkage TreeCluster. The default Cluster Picker threshold for Max-diameter clustering is 4.5%, [4], so we use this as our clustering threshold for Max-Diameter TreeCluster (both with and without the Clade constraint). For Sum-length TreeCluster (with and without the Clade constraint), we simply double the Max-diameter threshold and use 9%. In addition to using these default thresholds, we also test a wide range of thresholds for each transmission clustering method for robustness.

We measure cluster growth from year 8 to year 9 of the simulation and select the 1,000 highest-priority individuals, where individuals are prioritized in descending order of respective cluster growth. To measure the risk of a given individual *u*, we count the number of HIV transmission events *u*→*v* between years 9 and 10. To measure the effectiveness of a given clustering, we average the risk of the selected top 1,000 individuals and use this as a metric of how effective the clustering method is. Higher numbers imply the ability to prevent more transmissions by targeting a fixed (1,000) number of individuals and thus are desirable. As a control, we also show the mean number of transmissions per population, which is what a random selection of 1,000 individuals would give in expectation (we call this “expected” risk).

#### Application 3: Divide-and-conquer Multiple Sequence alignment

##### Algorithmic idea

Tree-based clustering has also been used for divide-and-conquer, a technique that has proved particularly useful for scaling existing methods for tree inference and multiple sequence alignment (MSA) to very large datasets [10, 43–47]. Divide-and-conquer methods first use some approach to build a quick-and-dirty estimate of the phylogeny and then divide the dataset into smaller sets using the phylogeny, such that sequences inside each subset are less diverse than the full set; given the subsets, an accurate (but often computationally demanding) method is run on the subsets to infer the MSA and/or the tree; finally, the results on the subsets are merged using various techniques. The accuracy of the output depends not only on the accuracy of the base method used on the subsets and the the merging method, but also, on the effectiveness of the method used to divide the tree into subsets [48].

Here, we specifically focus on MSA using divide-and-conquer. In particular, we focus on a method called PASTA that infers both MSAs and trees for ultra-large datasets (tested for up to 1,000,000 sequences). PASTA computes an initial alignment using HMMs implemented in HMMER [49] and an initial tree using FastTree-II [8]; then, it performs several iterations (default: 3) of the divide-and-conquer strategy described before using Mafft [50] for aligning subsets and using a combination of OPAL [51] and a technique using transitivity for merging subalignments. A tree is generated using FastTree-II at the end of each iteration, which is then used as the guide tree for the next iteration. The method has shown great accuracy on simulated and real data, especially in terms of tree accuracy, where it comes very close to the accuracy obtained using the true alignment, leaving little room for improvement. However, in terms of the alignment accuracy, it has substantial room for improvement on the most challenging datasets.

The divide-and-conquer of PASTA is based on the centroid-edge decomposition. Given the *guide tree* (available from the previous iteration), the decomposition is defined recursively: divide the tree into two halves, such that the two parts have equal size (or, are as close in size as possible). Then, recurse on each subtree, until there are no more than a given number of leaves (default: 200) in each subset.

##### Formulation as min-cut partitioning

The centroid edge decomposition involves cutting edges and includes a constraint defined on the subsets. However, it is defined procedurally and does not optimize any natural objective function. The min-cut partitioning can produce a decomposition that is similar to the centroid decomposition in its constraints but is different in outcome. Take the guide tree and set all edge weights to 1. Then solve our Sum-length min-cut partitioning problem with the threshold set to *α* = 2*m*-2; the result is a partition such that no cluster has more than *m* leaves and the number of subsets is minimized. Thus, this “max-size min-cut partitioning” is identical to centroid decomposition in its constraints, but also guarantees to find the minimum number of clusters.

##### Experiments

To evaluate how our new decomposition impacts PASTA, we compare the old version to a new version that uses the decomposition based on max-size min-cut partitioning, with other parameters (including maximum subset size) all kept fixed. We run both versions on two datasets both from the original PASTA paper: 10 replicates of a simulated RNAsim dataset with 10,000 leaves and a set of 19 real HomFam datasets with 10,099 to 93,681 protein sequences. The RNASim is based on a very complex model of RNA evolution. Here, the true alignment, known in simulations, is used as the reference. For HomFam, since the true alignment is not known, following previous papers, we rely on a very small number of seed sequences with a hand-curated reliable alignment as reference [45, 52]. In both cases, we measure alignment error using two standard metrics computed using FastSP [53]: SPFN (the percentage of homologies in the reference alignment not recovered in the estimated alignment) and SPFP (the percentage of homologies in the estimated alignment not present in the reference).

## Results

### Results for Application 1: OTU clustering

On the Greengenes dataset, as we change the threshold between 0.005 and 0.15, we get between 181, 574 and 10, 112 clusters (note that singletons are also counted). The cluster diversity has a non-linear relationship with the number of clusters; it drops quicker with higher thresholds where fewer clusters are formed (Figs. 3 and S1). Comparing the three objective functions that can be used in TreeCluster, we observe that Max-diameter and Sum-length have similar trends of cluster diversity scores whereas Single-linkage min-cut partitioning has substantially lower diversity compared to the other two methods (Fig. S1). This pattern is observed whether distances are computed using tree distances or sequence distances, but differences are larger for tree distance. Finally, note that even though tree distances are, as expected, larger than sequence distances (Fig. S2), the cluster diversity is *lower* when computed using tree distances, showing that clusters are tight in the phylogenetic space.

**Fig 3.**
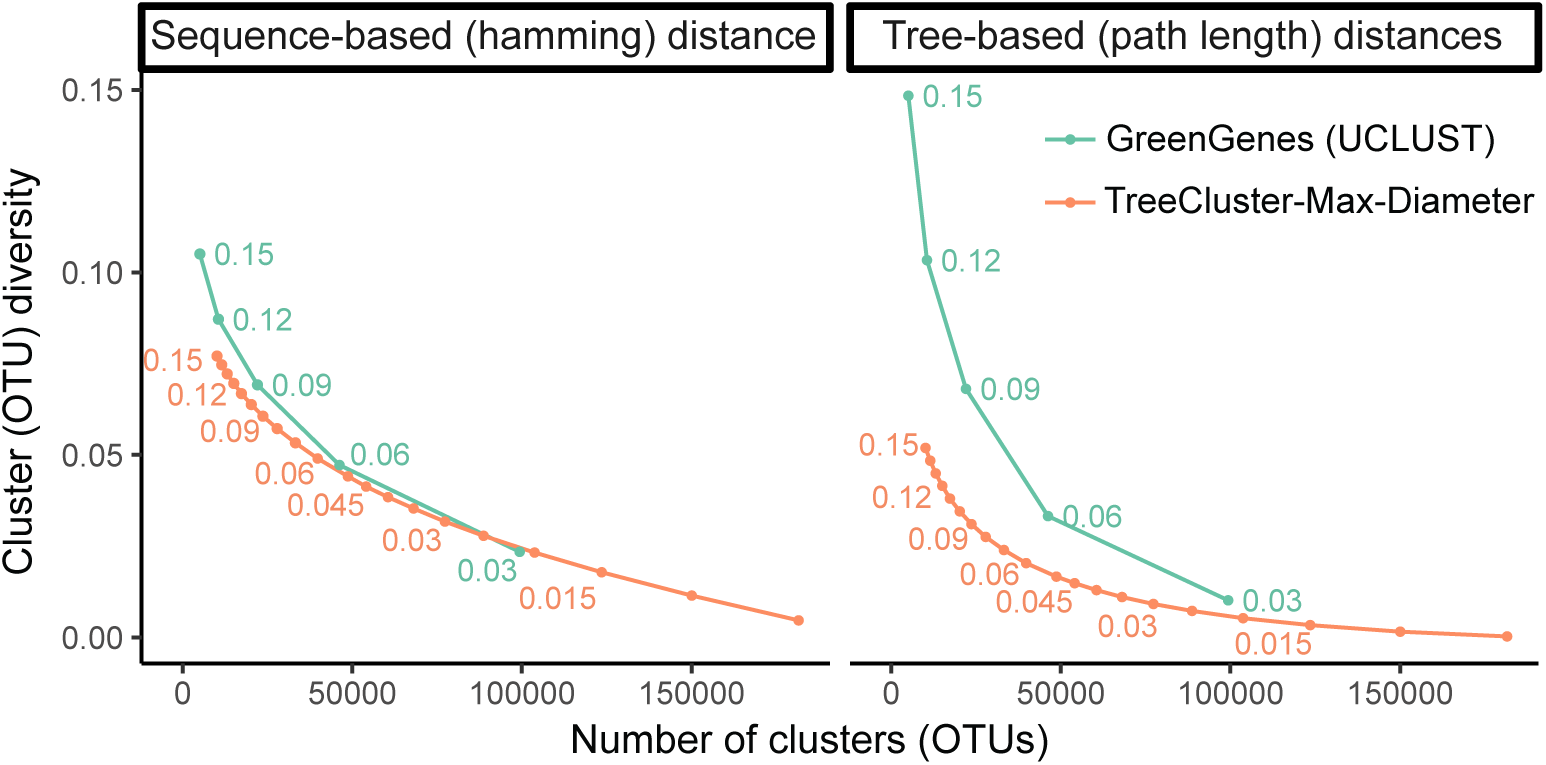
Clustering diversity, defined as weighted average pairwise distance within a cluster (Eq. 3) for Greengenes and TreeCluster versus the number of OTUs. For any number of OTUs (x-axis), a lower OTU diversity (y-axis) is preferrable. The threshold *α* is shown for all data points corresponding to GreenGeenes and for some points of TreeCluster. See Fig. S1 for comparison to other TreeCluster versions.

Compared to default Greengenes OTUs, defined using UCLUST, Max-diameter min-cut partitioning defines tighter clusters for tree-based scores (Fig. 3). When distances between sequences are measured in tree distance, cluster diversity score for Greengenes OTUs is substantially lower for all thresholds, and the gap is larger for higher thresholds. For example, the cluster diversity of Greengenes OTUs is three times higher than TreeCluster OTUs for *α* = 0.15. When distances between sequences are measured in hamming distance, Greengenes and TreeCluster perform similarly for low threshold values (e.g., note *α* = 0.03 for Greengenes, which is similar to *α* = 0.02 for TreeCluster in terms of the number of clusters). However, when the number of OTUs is reduced, remarkably, TreeCluster outperforms Greengenes OTUs by up to 1.4 folds (for *α* = 0.15). This is despite the fact that UCLUST is working based on sequence distances and TreeCluster is not.

Size of the largest cluster in Greengenes is larger compared to TreeCluster (Table 1). For example, for *α* = 0.09, both methods have similar number of clusters (22,090 and 23,631 for Greengenes and TreeCluster, respectively) but the size of largest cluster in Greengenes is three times that of TreeCluster (1,659 versus 540). On the other hand, for the same threshold value, the number of singleton clusters comprises 48% of all clusters for Greengenes whereas only 27% of the clusters are singletons for TreeCluster. Thus, GreenGenes has more clusters that are very small or very large, compared to TreeCluster.

**Table 1.**
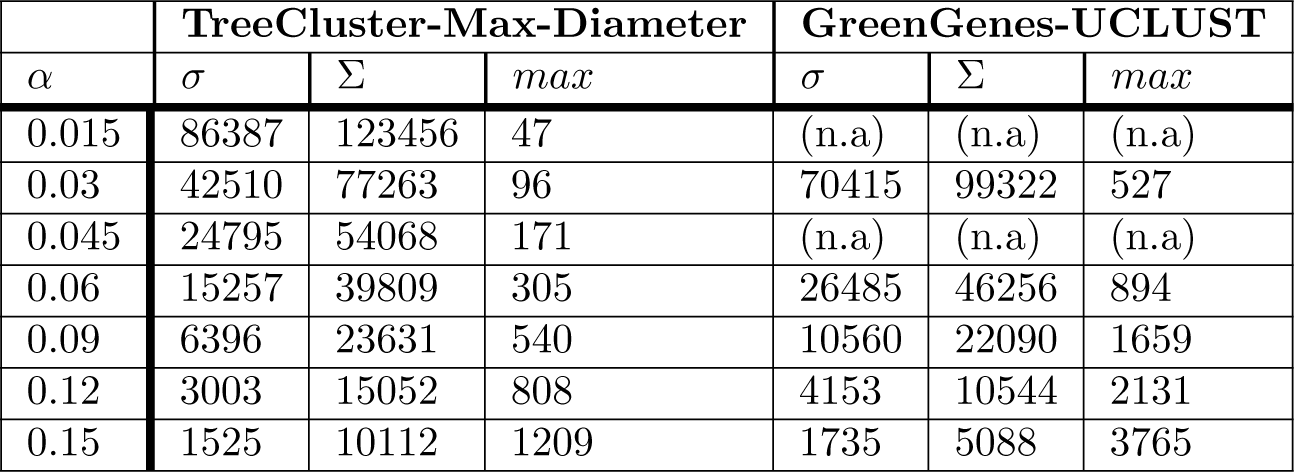
Number of singleton clusters (*σ*), total number of clusters (Σ), and maximum cluster size (*max*) for TreeCluster and GreenGenes on various threshold levels. In GreenGenes database, OTU definitions for threshold level *α* = 0.015 and *α* = 0.045 are not available.

### Results for Application 2: HIV dynamics

Comparing various versions of TreeCluster, regardless of the parameters that we vary, Sum-Branch Tree Cluster consistently outperforms the other clustering methods, and the inclusion of the Clade constraint has little impact on effectiveness (Fig. 4).

**Fig 4.**
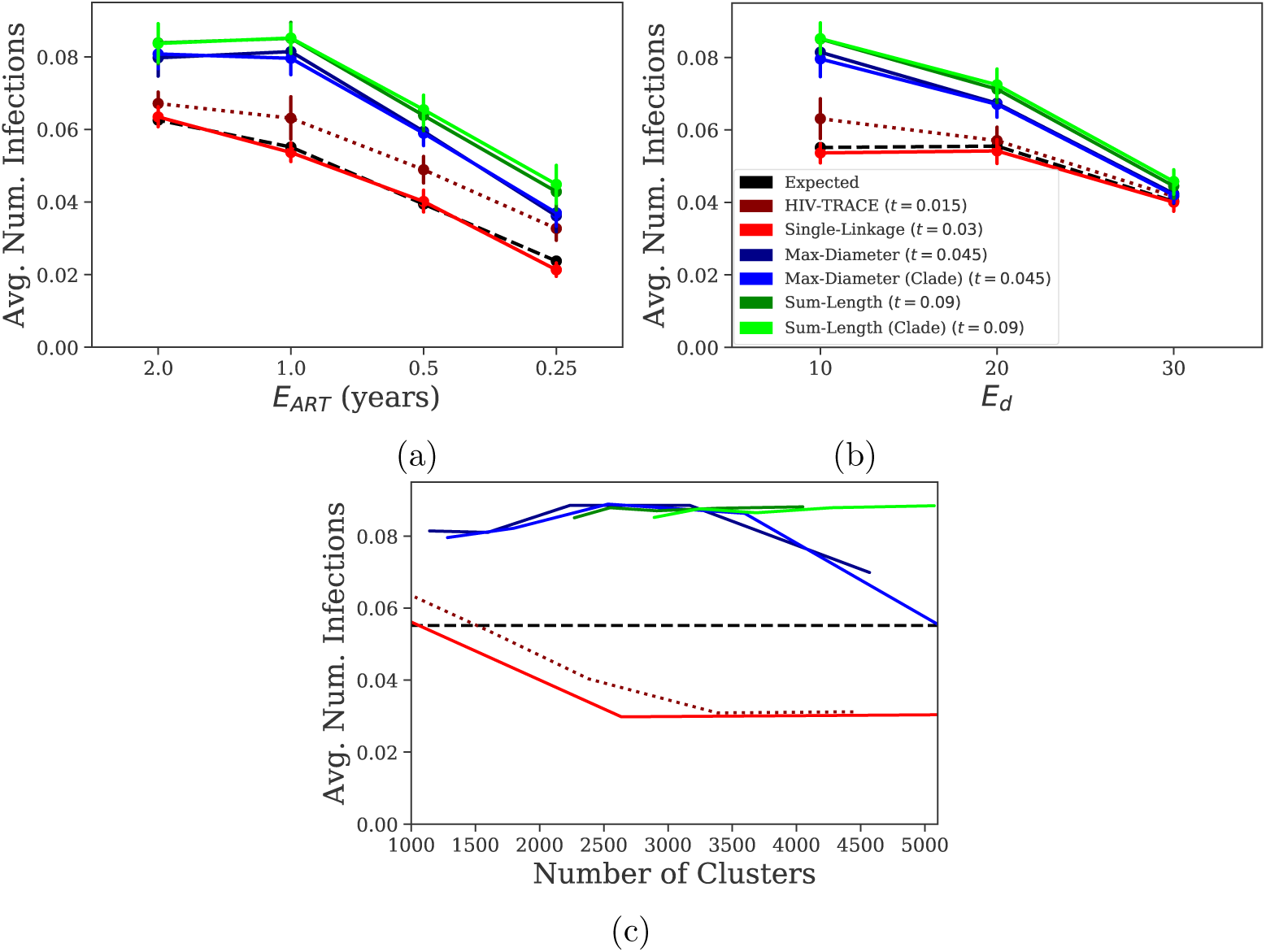
Effectiveness of transmission clustering, where effectiveness is measured as the average number of individuals infected by the selected 1,000 individuals. The horizontal axis depicts the expected time to begin ART (a), the expected degree (i.e., number of sexual contacts) for individuals in the contact network (b), and the number of clusters using various thresholds (c).

Compared to a random selection of individuals, the risk of selected individuals can be substantially higher; for example, with expected ART time set to 1 year, expected risk is 0.55 transmissions while the risk of top 1,000 individuals from Sum-length clusters is 0.85. In all the conditions, a close second to TreeCluster Sum-length, is TreeCluster Max-diameter. Other methods, however, are substantially less effective than these two modes of TreeCluster.

When varying expected time to begin ART and expected degree, Single-Linkage TreeCluster and HIV-TRACE consistently perform lower than the other approaches. Single-Linkage TreeCluster typically performing around the theoretical expectation of a random selection while HIV-TRACE performing slightly better (Fig. 4a-b). Recall that these two methods are conceptually similar. Moreover, these patterns are not simply due to the chosen thresholds. Even when we change the thresholds to control the number of clusters, Single-Linkage TreeCluster and HIV-TRACE consistently perform worse than expected by random selection (Fig. 4c). The effectiveness of other methods is maximized when they create between 2,000 to 5,000 clusters for Sum-length, or between 2,000 to 3,000 clusters for Max-Diameter.

### Results for Application 3: improving PASTA

When we replace centroid decomposition with max-size min-cut partitioning in PASTA, the alignment error reduces substantially for the RNASim dataset, but less so on the Homfam dataset (Fig. 5). On the RNASim data, mean SPFN drops from 0.12 to 0.10, which corresponds to a 17% reduction in error. These drops are consistent across replicates and are substantial given the fact that the only change in PASTA was to replace its decomposition step with our new clustering algorithm, keeping the rest of the complex pipeline unchanged. In particular, the method to align subsets, to merge alignments, and to infer trees, were all kept fixed. On the HomFam dataset, too, errors decreased, but the reductions were not substantial (Fig. 5b). Based on these results, we have now changed PASTA to use max-size min-cut partitioning by default.

**Fig 5.**
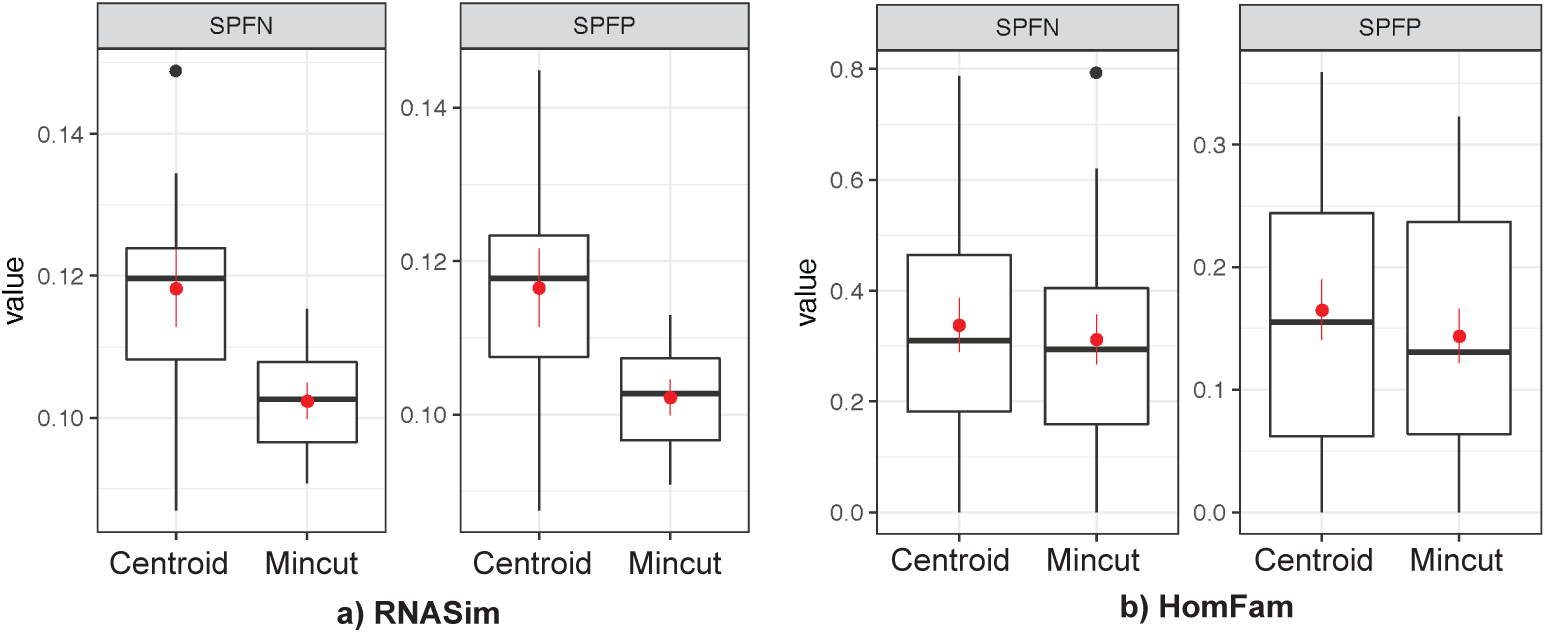
Alignment error for PASTA using the old (centroid) and the new (mincut) decompositions. We show Sum of Pairs False Negative (SPFN) and Sum of Pairs False Psotive (SPFP) computed using FastSP [53] over two datasetes: Simulated RNASim dataset (10 replicates) with 10 replicates and biological HomFam dataset (19 largest families; all 20 largest, except “rhv” omitted due to the warning on the Pfam website). We show boxplots in addition to mean (red dot) and standard error (red error bars).

## Discussion

Several theoretical and practical issues should be further discussed.

### Mean-diameter min-cut partitioning

Some of the existing methods, such as ClusterPicker [4], can define their constraints based on mean pairwise distance between nodes. Similar to those, we can define a variation of the min-cut partitioning problem where 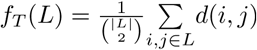. Unfortunately, this “mean-diameter” min-cut partitioning problem can be solved in linear time using our greedy algorithm only if we also have clade constraints (Algorithm 4 in Appendix B). As demonstrated by the counter example given in Fig. S3, the greedy algorithm fails if we do not have clade constraints. More generally, the use of mean as function *f*_*T*_ (.) creates additional complexity and we conjecture the problem will not be solvable in linear time. Whether mean diameter is in fact a reasonable criteria is not clear. For example, it is possible that the mean diameter of a cluster is below the threshold while the mean diameter of clusters embedded in that cluster are not; such scenarios may not make sense for downstream applications.

### Set of optimal solutions

It is possible that multiple distinct partitions with equal number of clusters are all optimal solutions to any of our min-cut partitioning problems. Moreover, as the example given in Figure 6 shows, the number of optimal solutions can be exponential with respect to number of leaves in a binary phylogenetic tree. This observation renders listing all optimal solutions of diminished interest since there could be too many of them. However, finding a way to summarize all partitions may have practical utility. We do not currently have such a summarization approach. However, as shown in Lemma S1 of Appendix B, although the optimal solution space is potentially exponentially large, one can easily determine the set of all edges that could appear in any of the optimal solutions. Thus, we could find absolutely unbreakable edges that will not be cut in any optimal clustering of the data.

**Fig 6.**
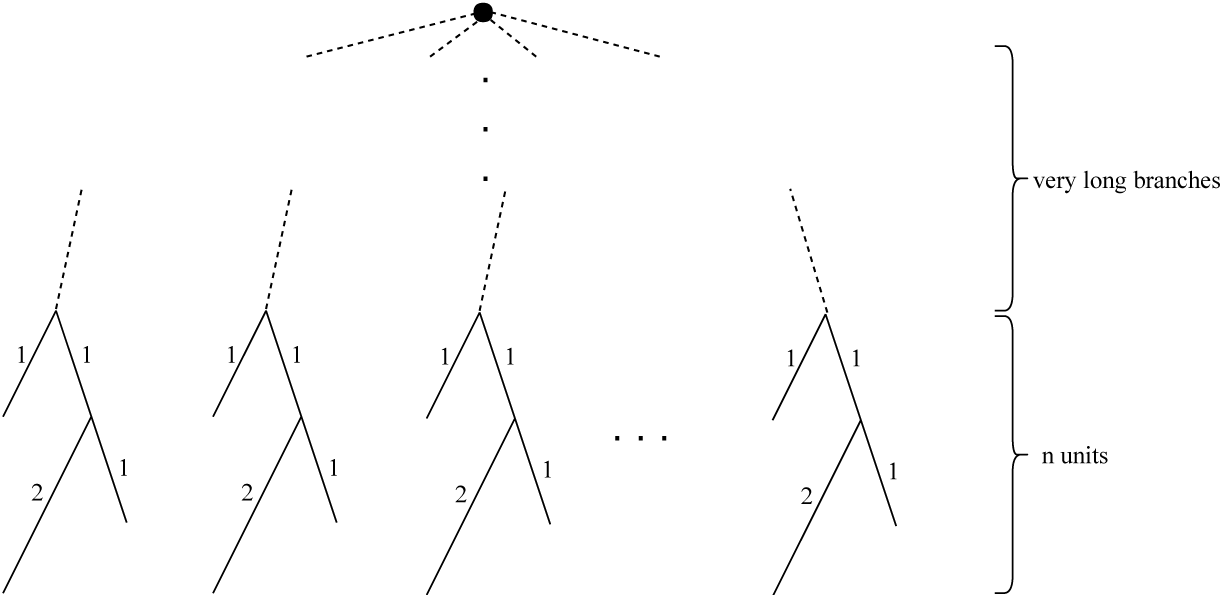
An example showing that number of minimal clusterings under diameter threshold can be exponential of number of leaves. When threshold is equal to 3.5, each unit has to be split into two clusters and there are three equally legitimate way of splitting. The minimum of clusters is therefore 2*n*. Total number of distinct optimal solutions is 3^*n*^ whereas there are 3*n* leaves.

### Choice of criterion

Among the three methods that we discussed, we observed that Max-diameter and Sum-diameter are consistently better than the Single-linkage. This observation makes intuitive sense. Single-linkage can increase the diversity within a cluster simply due to the transitive nature of its criterion. Thus, a very heterogeneous dataset may still be collapsed into one cluster, simply due to transitivity. Our desire to solve the Single-linkage problem was driven by the fact that a similar concept is used in HIV-TRACE, arguably the most widely used HIV clustering method. However, we did not detect any advantage in this type of clustering compared to Max-diameter or Sum-length; thus, our recommendation is to use these two criteria instead. Between the two, Max-diameter has the advantage that its *α* threshold is easier to interpret.

### Centroid node

OTU picking methods, in addition to clustering, also choose a centroid node per cluster. Our clustering approach is centroid-free. However, if a representative is needed, many natural choices are available. For example, one can in linear time find the midpoint of a cluster or its balance point [15]; then, the leaf closest to the midpoint or balance point can be used as the representative. Alternatively, as some recent papers argue [54], constructing and using a consensus sequence (or perhaps even a reconstructed ancestral sequence) may be preferable to using one of the given sequences as the centroid.

### Running time

We focused on comparing effectiveness of TreeCluster to other methods, but we note that its running time also compares favorably to other clustering methods (once the tree is inferred). For example, on a real HIV dataset, we ran HIV-TRACE, Cluster Picker, and TreeCluster for subsets of the data ranging from 100 to 5,000 sequences (Fig. 7). Even on the largest dataset, the running time of TreeCluster did not exceed 2 seconds. In contrast, the sequence-based HIV-Trace required close to a minute, (which is still quite fast) but Cluster Picker needed more than an hour. Even on the Greengenes dataset with more than 200,000 leaves, TreeCluster performed clustering in only 30 seconds. The high speed of TreeCluster makes it possible to quickly scan through a set of *α* thresholds to study its impact on the outcomes of downstream applications.

**Fig 7.**
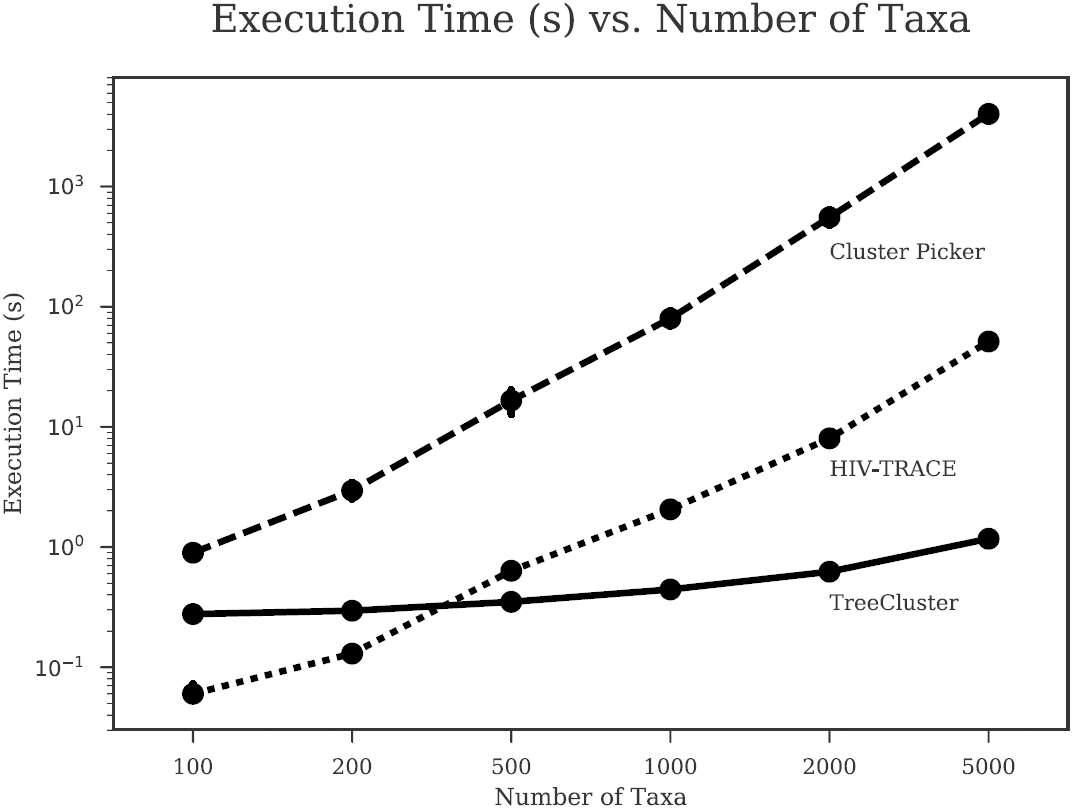
Execution times of Cluster Picker, HIV-TRACE, and TreeCluster in log-scale. Execution times (in seconds) are shown for each tool for various values of *n* sequences, with 10 replicates for each *n*. The full dataset was obtained by downloading all HIV-1 subtype B *pol* sequences (HXB2 coordinates 2,253 to 3,549) from the Los Alamos National Laboratory (LANL) database. All programs were run on a CentOS 5.8 machine with an Intel Xeon X7560 2.27 GHz CPU.

We note that these numbers do not include the time spent for inferring the tree, which should also be considered if the tree is not already available. For example, based on previous studies, MSA and tree inference on datasets with 10,000 sequences can take close to an hour using PASTA and 12 CPUs. Around a third of this time is spent on tree inference (e.g., see Figure 4 of [45]) and the rest is spent on the estimating alignment, which is also needed by most alternative clustering methods.

## Conclusion

We introduce TreeCluster, a method that can cluster sequences at the tips of a phylogenetic tree using several optimization problems. We showed that our liner time algorithms can be used in several downstream applications, including OTU clustering, HIV transmission clustering, and divide-and-conquer alignment. Using the tree to build the cluster increases their internal consistency and improves downstream analyses.

## Acknowledgments

Authors thank Xingfan Jia for useful discussions.

## A Supporting information

**Fig S1.**
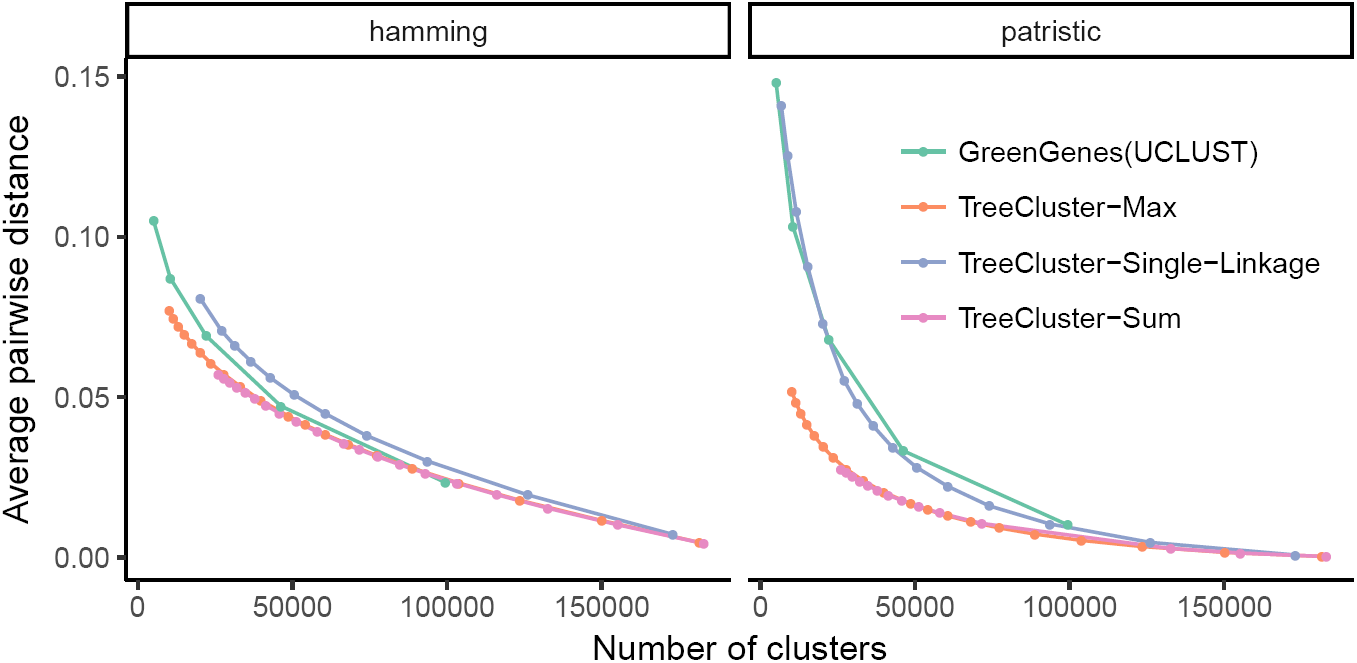
Comparison of various TreeCluster options and Greengenes. Clustering quality of Greengenes and various options of TreeCluster, where quality is measured as average pairwise distance within a cluster (the lower the better). The horizontal axis shows the number of clusters for a given method and a threshold value. TreeCluster OTUs based on Max-diameter and Sum-length options outperform Single-linkage option as well as Greengenes OTUs. Computation of Hamming distance based cluster diversity for *α* ≥ 0.7 did not complete within 24 hours and had to be terminated.

**Fig S2.**
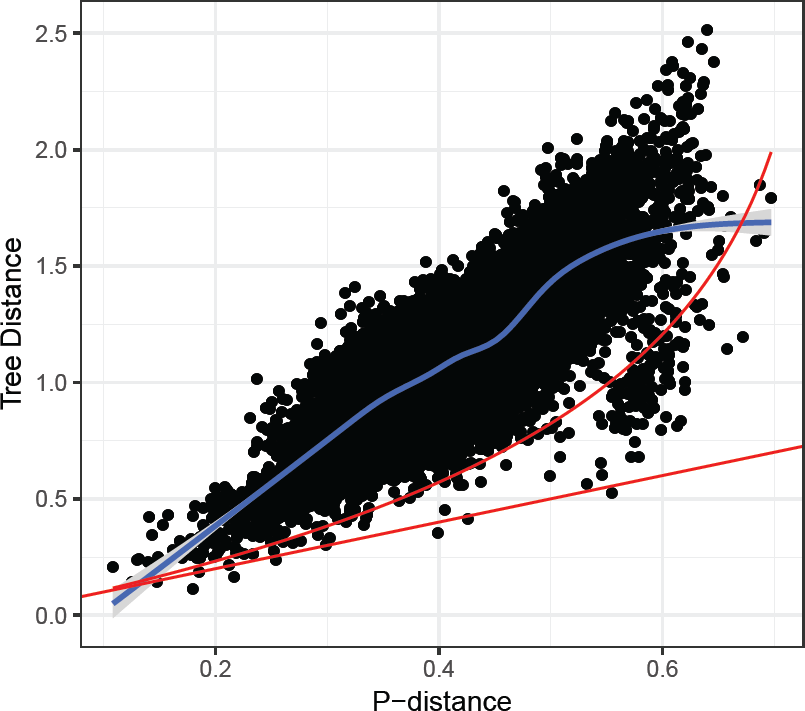
Tree distance versus hamming distance. On 16S data, the relationship between tree distances and hamming distances cannot be established using Jukes-Cantor formula (red curve).

**Fig S3.**
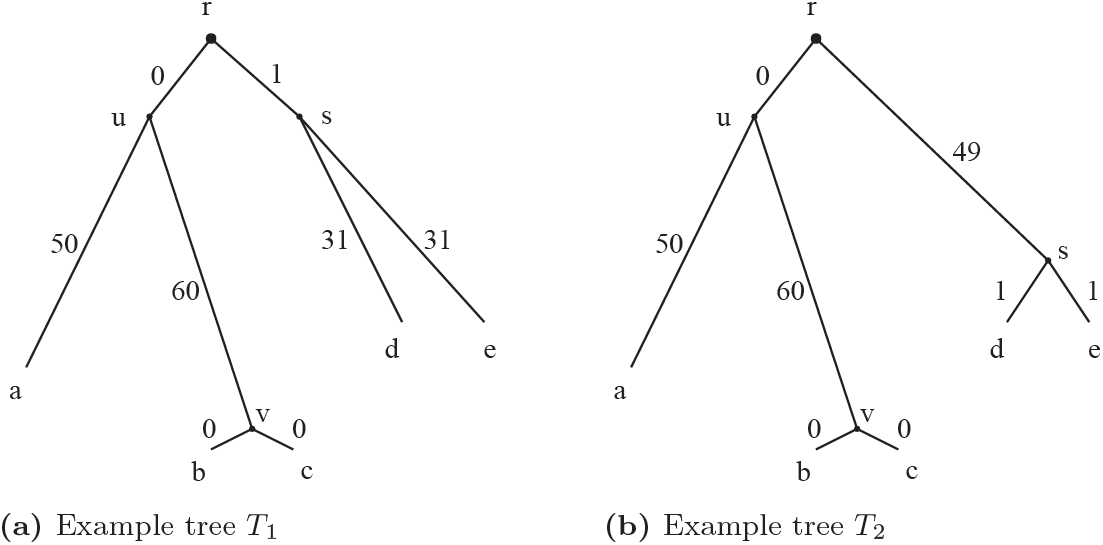
An example showing that mean-diameter min-cut partitioning is not conforming locality when *α* = 72, thus cannot be solved by a greedy algorithm analogous to Alg. 1. When a greedy algorithm is at the stage where it processes *u*, it makes the decision for cutting its children edges (*u, v*) and (*u, a*) based on the information available at the subtree rooted by *u*. When *α* = 72, *T*_1_ and *T*_2_ require different cut-sets (*{*(*u, v*)} and {(*u, a*)} respectively) for the optimal Mean-diameter partitioning despite the fact that the subtree rooted by *u* remains unchanged in *T*_1_ and *T*_2_.

## B Proofs and supplementary algorithms

### B.1 Linear solution for Sum-length min-cut partitioning problem

#### Algorithm 3: Linear solution for Sum-length min-cut partitioning

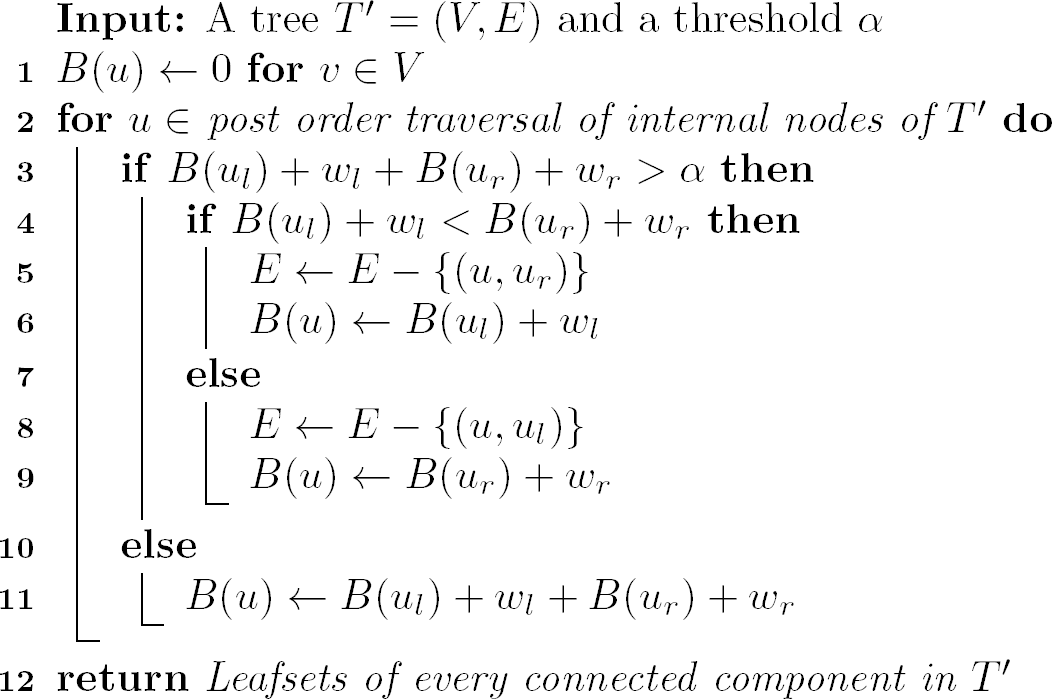

We now show the Algorithm 3 correct. Let *A*(*u*) be the minimum number of clusters under *U* all with a diameter less than *α*; i.e. *A*(*r*) is the objective function.

#### Theorem S1.

*Algorithm 1 computes a clustering with minimum A*(*r*) *for rooted tree T* ′. *In addition, among all possible such clusterings, the algorithm picks the solution with minimum B*(*r*).

*Proof*. The proof uses induction. The base case for the induction is the simple rooted tree with root *u* and two leaves *u*_*l*_ and *u*_*r*_. If *w*_*l*_ + *w*_*r*_ > *α* the algorithm cuts the longer branch whereas if *w*_*l*_ + *w*_*r*_ ≤ *α* no branch is cut. In both cases, the theorem holds.

When *B*(*u*_*l*_) + *w*_*l*_ + *B*(*u*_*r*_) + *w*_*r*_≤*α*, we have *A*(*u*) = *A*(*u*_*l*_) + *A*(*u*_*r*_)-1, which is the minimum possible by inductive hypothesis and the fact that the number of clusters cannot go down by more than one on node *u*. Also, *B*(*u*) is optimal by construction.

When *B*(*u*_*l*_) + *w*_*l*_ + *B*(*u*_*r*_) + *w*_*r*_ > *α*, without loss of generality, assume that *B*(*u*_*l*_) + *w*_*l*_ ≥ *B*(*u*_*r*_) + *w*_*r*_ and thus, the algorithm cuts the (*u, u*_*l*_) branch, getting *A*(*u*) = *A*(*u*_*l*_) + *A*(*u*_*r*_) and *B*(*u*) = *B*(*u*_*r*_) + *w*_*r*_. Note that *A*′(*u*) < *A*(*u*) is only possible if *A*′(*u*_*l*_) = *A*(*u*_*l*_) and *A*′(*u*_*r*_) = *A*(*u*_*r*_) and we do not cut any branch at *u* in the alternative clustering. However, this scenario is *not* possible because

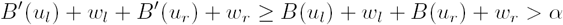

where the first inequality follows from the inductive hypothesis and the final inequality shows that we will have to cut a branch in any alternative setting. Finally, we need to show that an alternative solution with *A*′(*u*) = *A*(*u*) but *B*′(*u*) < *B*(*u*) is not possible. The inequality requires that either *B*′(*u*_*l*_) < *B*(*u*_*l*_) or *B*′(*u*_*r*_) < *B*(*u*_*r*_). First, consider the *B*′(*u*_*l*_) < *B*(*u*_*l*_) case, which is possible only if *A*′(*u*_*l*_) = *A*(*u*_*l*_) + 1. Note that *A*′(*u*) = *A*(*u*) requires *A*′(*u*_*r*_) = *A*(*u*_*r*_) (and thus *B*′(*u*_*r*_) = *B*(*u*_*r*_)) and that *B*′(*u*_*l*_) + *w*_*l*_ + *B*(*u*_*r*_) + *w*_*r*_ < *α*, which is possible. Under this condition, we find:

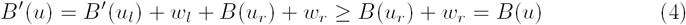

If instead *B*′(*u*_*r*_) < *B*(*u*_*r*_), similar conditions can be written, resulting in

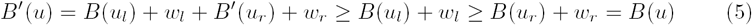

Thus, *A*(*u*) and *B*(*u*) are optimal when *B*(*u*_*l*_) + *w*_*l*_ + *B*(*u*_*r*_) + *w*_*r*_ > *α*.

#### Corollary S1

*Let C*′ *be the cut set obtained by running Alg. 1 on any arbitrary rooting T* ′ *of unrooted tree T. C*′ *optimally solves the Max-diameter min-cut partitioning problem*.

*Proof*. Let *r*_*r*_ and *r*_*l*_ denote the right and the left child of the root of *T* ′. Every edge in *T* can be mapped to *T* ′ except the edge (*r*_*right*_, *r*_*left*_), from which we define a mapping to (*r, r*_*right*_) (w.l.o.g). Using this mapping, the optimal clustering (i.e. optimal cut-set) on *T* can be translated to an alternative max diameter min-cut partitioning on *T* ′. However, by Theorem 1, *A*(*r*) is optimal and cannot be improved by any alternative partitioning. Since any admissible clustering on *T* ′ is also admissible on *T*, Alg. 1 minimizes *q*.

### B.2 Proofs for Single-linkage min-cut partitioning problem

*Proof of Proposition 1*. (⇐) If *d*(*a, b*) ≤ *α* but *a* and *b* are in distinct clusters *L*_*a*_, *L*_*b*_ respectively, *N* can be reduced by one by simply merging *L*_*a*_ and *L*_*b*_. *f*_*T*_ (*L*_*a*_ ∪ *L*_*b*_) ≤ *α* is satisfied if for any split of *L*_*a*_ ∪ *L*_*b*_, there exists a pair of leaves that are from distinct splits and are within *α* threshold. For any pair of non-empty sets *S* and *S*′ that satisfy *S ⊂ L*_*a*_ and *S*′ *⊂ L*_*b*_, we have 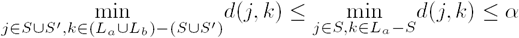 and 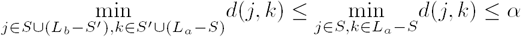. On the other hand, 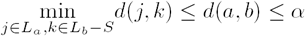. This concludes that for *L* = *L*_*a*_ ∪ *L*_*b*_, *f*_*T*_ (*L*) ≤ *α* is satisfied. *L*_*a*_ and *L*_*b*_ can still be merged if the chain ℋ described above exists. It is trivial to show that there is a link ⟨*c*_*i*_, *c*_*i*+1_⟩ in ℋ such that *c*_*i*_ *∈ L*_*a*_ and *c*_*i*+1_≠*L*_*a*_. Using the argument above, we can iterate over *ℋ* and keep merging clusters (and decrease N) every time we see such a link until we finally merge *L*_*a*_ with *L*_*b*_.

(⇒) We describe a procedure to compute the chain *ℋ*. If *a* and *b* in the same cluster *L*, 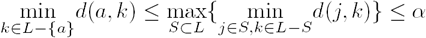 holds, implying that there is a leaf *c*_1_ in set *L - {a}* such that *d*(*a, c*_1_) ≤ *α*. If *c*_1_ = *b*, theorem follows. If *c*_1_≠*b*, we union *a* and *c*_1_, call the union set *L*_*a*_, and add the link *a* → *c*_1_ to ℋ′. Iteratively, we find the pair ⟨*j, k*⟩ that yields to 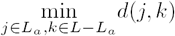, add the link *j* → *k* to ℋ′, and add *k* to *L*_*a*_ until we finally add *b* to *L*_*a*_. The elements forming the path between *a*, and *b* in ℋ′, which can be computed using depth-first-search, constitute a valid chain *ℋ*.

### B.3 Average-clade clustering

#### Algorithm 4 Average Diameter Clade Average diameter clade min-cut partitioning

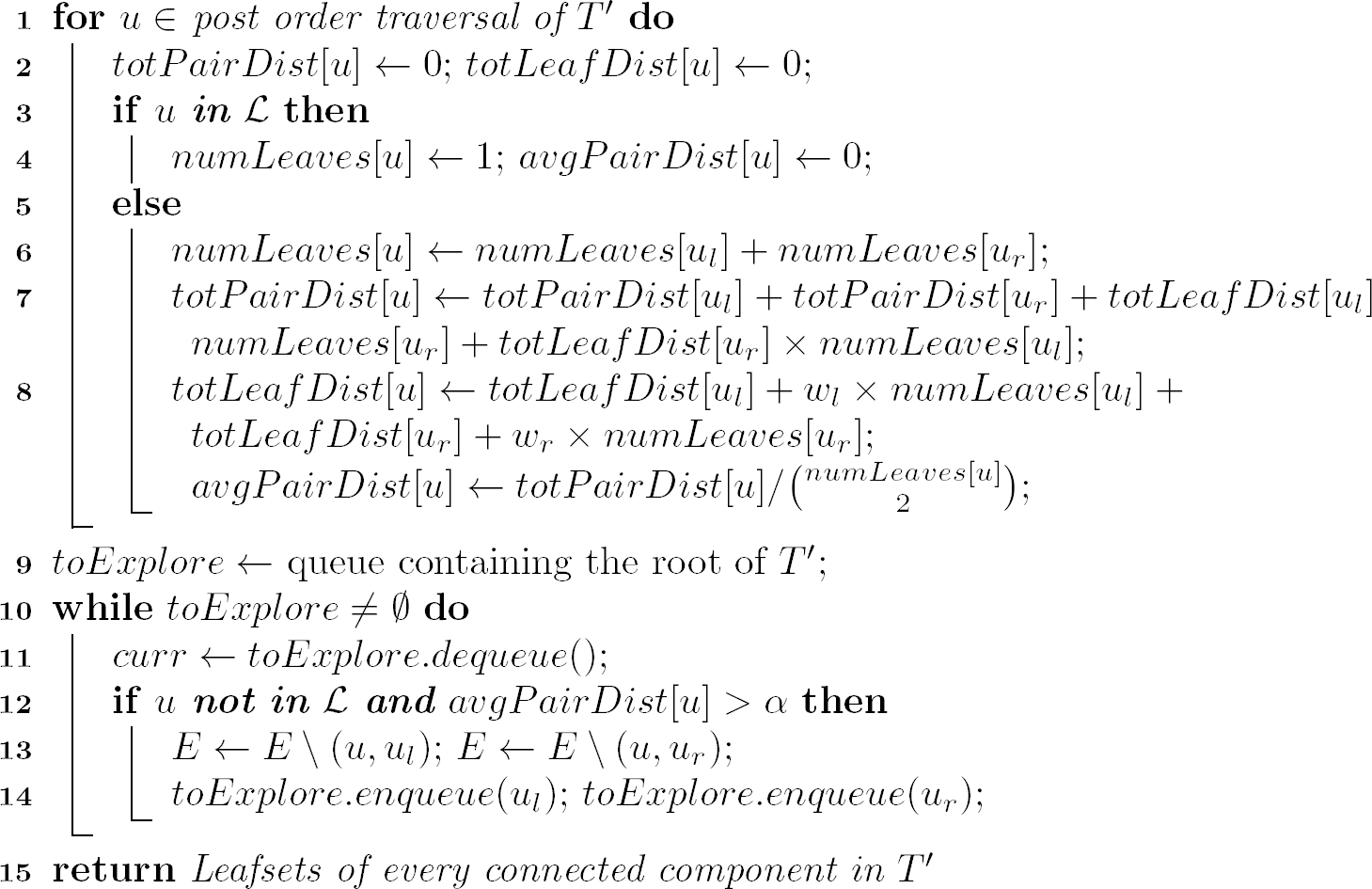

### B.4 All optimal solutions

#### Lemma S1

*Let {e*_1_, *e*_2_,…, *e*_*m*_*} be the set of edges in an unrooted tree T. Consider the following algorithm: root T at e*_*j*_ *and run Max algorithm 𝒮*_*j*_, *and let* _*j*_ *be the set of edges cut by the algorithm in this run. Any optimal clustering for T has to draw its cut set from* 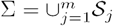.

*Proof*. The proof is by contradiction. Assume there is an optimal cut set 𝒮′ that does contain an edge *e*_*i*_ that *e*_*i*_ ∉ Σ. Consider the *T* rooted at *e*_*i*_. We call root of this tree as *v*, immediate left and right branches of *v* as *e*_*l*_ and *e*_*r*_, and left and right child nodes of *v* as *v*_*l*_ and *v*_*r*_. Note that *e*_*l*_ and *e*_*r*_ correspond to *e*_*i*_ in *T*. Because of our assumption, *e*_*l*_ ∉ *𝒮*_*j*_ and *e*_*r*_ ∉ *𝒮*_*j*_. When *e*_*i*_ is removed from *T*, two new trees form, called *T*_*left*_ (the one containing the node *v*_*l*_) and *T*_*right*_ (the one containing the node *v*_*r*_). If *p* number of cuts in 𝒮′ are in *T*_*right*_, and *q* number of cuts in 𝒮′ are in *T*_*left*_, |𝒮′| equals *a* + *b* + 1. Number of cuts in 𝒮′ and *𝒮*_*j*_ are equal and *e*_*l*_ and *e*_*r*_ is not cut, which implies that either the tree rooted by *v*_*l*_ or *v*_*r*_ has an alternative clustering with one less cut. By the design of the *Max* algorithm, if this was the case, algorithm would choose the alternative cut.

## Footnotes

1 In fact, in our quest to design tree-based algorithms, we reinvented these same algorithms only to later find out that they have been previously described.

